# Polaris: Polarization of ancestral and derived polymorphic alleles for inferences of extended haplotype homozygosity in human populations

**DOI:** 10.1101/2024.12.06.627098

**Authors:** Alessandro Lisi, Michael C. Campbell

**Affiliations:** Department of Biological Sciences (Human and Evolutionary Biology Section), University of Southern California, 3616 Trousdale Parkway, Los Angeles, CA 90089

**Keywords:** ancestral alleles, derived alleles, natural selection, integrated haplotype score (*i*HS), extended haplotype homozygosity (EHH)

## Abstract

**Summary:** Statistical methods that measure the extent of haplotype homozygosity on chromosomes have been highly informative for identifying episodes of recent selection. For example, the integrated haplotype score (*i*HS) and the extended haplotype homozygosity (EHH) statistics detect long-range haplotype structure around derived and ancestral alleles indicative of classic and soft selective sweeps, respectively. However, to our knowledge, there are currently no publicly available methods that classify ancestral and derived alleles in genomic datasets for the purpose of quantifying the extent of haplotype homozygosity. Here, we introduce the Polaris package, which polarizes chromosomal variants into ancestral and derived alleles and creates corresponding genetic maps for analysis by selscan and HaploSweep, two versatile haplotype-based programs that perform scans for selection. With the input files generated by Polaris, selscan and/or HaploSweep can produce the appropriate sign (either positive or negative) to outlier *i*HS statistics, enabling users to distinguish between selection on derived or ancestral alleles. In addition, Polaris can convert the numerical output of these analyses into graphical representations of selective sweeps, increasing the functionality of our software.

**Results:** To demonstrate the utility of our approach, we applied the Polaris package to Chromosome 2 in the European Finnish population from the 1000 Genomes Project. More specifically, we examined the regulatory region in intron 13 of *MCM6* associated with lactase persistence (i.e., the ability to digest the lactose sugar present in fresh milk), a region of intense interest to human evolutionary geneticists. Our analyses showed that the derived T_-13910_ allele (a known enhancer for lactase expression), sits on an extended haplotype background in the Finnish consistent with a classic selective sweep model as determined by *i*HS and EHH statistics calculated by selscan and HaploSweep. Importantly, we were able to immediately identify this target allele under selection based on the information generated by our software. We also explored outlier statistics across Chromosome 2 in two distinct datasets: i) one containing polarized alleles generated with Polaris and ii) the other containing unpolarized alleles in the original phased vcf file. Here, we found a significant excess of outlier statistics (*P* < 0.00001) in the unpolarized dataset, raising the possibility that a subset of these ‘hits’ of selection on Chromosome 2 may be false positives. Overall, Polaris is a versatile package that enables users to efficiently explore, interpret, and report signals of recent selection in genomic datasets.

**Availability and implementation:** The Polaris package is free and open source on GitHub (https://github.com/alisi1989/Polaris) and on DropBox (https://www.dropbox.com/scl/fo/mlxizft5267vem9u62qkn/AAnM0qX923zPzQBlPX8iteM?rlkey=uezrp4t2waffpj0nmo1evr320&e=1&st=jaodccws&dl=0)

## 1 Introduction

Positive natural selection is the process by which advantageous alleles rapidly rise to high frequency within populations, and it has played a critical role in the recent evolution of modern humans (Sabeti *et al*. 2002; Liu *et al*. 2013; Zhao *et al*. 2017; Harris, Garud and DeGiorgio 2018). To date, several haplotype-based statistical tests have been developed to detect episodes of positive selection (Sabeti *et al*. 2002, 2007; Hanchard *et al*. 2006; Voight *et al*. 2006; Wang *et al*. 2006; Zhang *et al*. 2006; Cai *et al*. 2011; Han and Abney 2013; Ferrer-Admetlla *et al*. 2014; Szpiech and Hernandez 2014; Zhao *et al*. 2024). Among them, the most widely used statistics are the integrated haplotype score (*i*HS; (Voight *et al*. 2006) and the extended haplotype homozygosity (EHH; (Sabeti *et al*. 2002), which measure the extent of haplotype homozygosity around adaptive alleles.

To calculate the *i*HS and EHH statistics, phased haplotypes, associated genetic maps, and in some cases the classification of polymorphisms as either ancestral or derived are typically required (depending on the software). Furthermore, the polarization of alleles can also provide a more nuanced understanding of past selective events. For example, large negative *i*HS statistics—as described in Voight et al. (Voight *et al*. 2006)—indicate that extensive haplotype homozygosity around derived alleles congruent with a classic selective sweep model. Under this scenario of selection, a beneficial allele appears on a single haplotype and rises to high population frequency (Lange and Pool 2016; Booker, Jackson and Keightley 2017; Rees, Castellano and Andrés 2020); conversely, large positive *i*HS statistics denote long-range haplotype homozygosity around ancestral alleles indicative of selection on standing variation (an example of a soft selective sweep)(Voight *et al*. 2006). In this circumstance, ancestral alleles previously segregating neutrally become selectively advantageous in a changing environment and increase in prevalence within a population (Pritchard, Pickrell and Coop 2010; Lange and Pool 2016; Rees, Castellano and Andrés 2020). Given the existence of these different modes of selection, large positive and negative *i*HS statistics are of particular interest in human evolutionary studies. However, there is currently no formalized method for categorizing ancestral and derived alleles in phased genomic datasets for the purpose of calculating these haplotype-based statistics. As a result, studies often ignore the sign (positive or negative) associated with *i*HS and report the absolute values instead. One consequence of this approach is the inability to immediately discern which allele at a given locus is adaptive as well as the possible mode of selection acting on this variation (i.e., a classic selective sweep or selection on standing variation).

To address this problem, we developed Polaris to recode phased alleles as either ancestral or derived along chromosomes and generate corresponding genetic maps for analysis by two state-of-the-art programs, selscan and HaploSweep (Szpiech and Hernandez 2014; Zhao *et al*. 2024), which scan for selection with a range of statistics, including *i*HS and EHH. These haplotype-based programs, using the input files generated by Polaris, can provide the appropriate sign (either positive or negative) to outlier statistics, enabling users to immediately distinguish between selection on derived or ancestral alleles. In addition, the Polaris package has the capability to visualize the numerical results of these analyses, facilitating the discovery, evaluation, and reporting of signals of recent selection.

## 2 Implementation

With the increased availability of whole genome sequencing and genomic genotyping data, researchers now have more power to identify distortions in haplotype structure following a selective sweep (Harris, Garud and DeGiorgio 2018). While many studies of positive selection have focused on detecting classic selective sweeps (Sabeti *et al*. 2002, 2006; Voight *et al*. 2006; Enard, Messer and Petrov 2014; Uricchio *et al*. 2019), data have indicated that other forms of selection, such as soft sweeps (specifically, selection on standing variation), have also played a key role in human adaptation and evolution (Hernandez *et al*. 2011; Messer and Petrov 2013; Schrider and Kern 2017).

Polaris was developed to classify alleles as ancestral or derived along chromosomes and create associated genetic maps for downstream haplotype-based analyses by selscan and HaploSweep—two programs that calculate *i*HS and EHH statistics among others. After categorizing variants as either ancestral or derived, selscan and HaploSweep will provide the appropriate sign (either positive or negative) to these statistics, thereby discriminating which alleles have likely undergone a selective sweep. Polaris can also convert the tabular output of these analyses into annotated Manhattan and/or color-coded line plots, broadening its versatility (**Fig. 1**).

**Figure 1.**
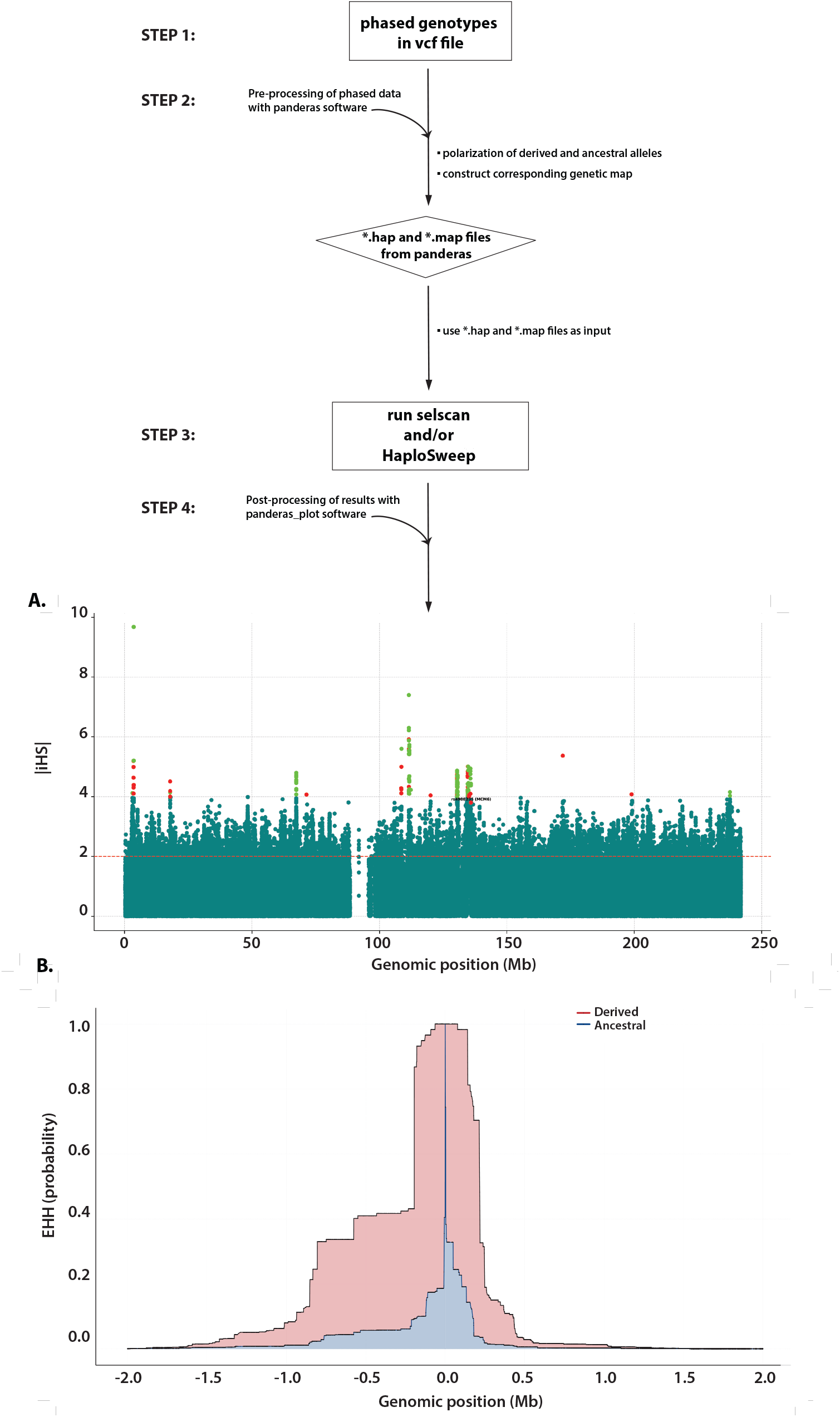
Workflow of the Polaris package. Polaris consists of two distinct programs: (i) Panderas and (ii) Panderas_Plots that run in a command-line Terminal. The input for the Panderas software consists of phased bi-allelic genotypes in a variant call format file without missing rs identifiers in the ID column (**Step 1**). The Panderas software recodes alleles in a phased vcf file as either ancestral or derived and constructs a corresponding genetic map. Then, this software generates two formatted files, *.hap and *.map, for direct use by selscan and/or HaploSweep (**Steps 2 and 3**). The Panderas_Plots software can visualize and annotate the tabular output of haplotype-based analyses as Manhattan and/or color-coded line plots (**Step 4**). For example, **Panel A** shows a Manhattan plot of *i*HS statistics calculated by selscan (Szpiech and Hernandez 2014) in the Finnish population from the 1000 Genomes Project. The dashed horizontal line in this plot indicates the threshold for outlier *i*HS statistics as specified by the user. We also highlighted the derived T_-13910_ allele associated with lactase persistence with a red dot and its corresponding rs identifier, rs4988235. The green and other red dots denote even more extreme *i*HS statistics. More explicitly, *i*HS > 4 (red dots) and *i*HS < -4 (green dots) denote selection on derived alleles (indicating a classic selective sweep) and selection on standing variation (indicating a soft selective sweep), respectively. **Panel B** (EHH) shows the decay of haplotype homozygosity with increasing distance from a core site (represented here by rs4988235). In this EHH line graph calculated by selscan (Szpiech and Hernandez 2014), the negative and positive numbers on the x-axis show the distance in Mb upstream and downstream from the core site (*rs*4988235) on the forward strand; the y-axis is the probability that two chromosomes are homozygous at all SNPs for the interval from the core site to distance x. Lastly, the blue line represents the decay of homozygosity of chromosomes carrying the ancestral allele at the core, while the red line signifies the decay of homozygosity on chromosomes with the derived allele at the core site.

### 2.1 Polarization of ancestral and derived alleles along chromosomes

The Polaris package consists of two distinct programs written in the C-Python language: (i) Panderas and (ii) Panderas_Plots.

#### 2.1.1 Step 1: Preparation of phased vcf file

The input for Panderas should contain phased bi-allelic genotypes in a variant call format (vcf) file. The task of inferring haplotypes from genotype or sequence data (also known as phasing) can be accomplished with a number of publicly available programs, such as Beagle or SHAPEIT (Delaneau, Marchini and Zagury 2011; Browning *et al*. 2021). Notably, all variants must have an rs identifier in the ID column of the vcf file. In cases where the vcf contains missing rs identifiers (e.g. “.”), users can apply a custom Python script provided in our GitHub repository to create unique identifiers in the ID column.

#### 2.1.2 Step 2: Recode phased alleles in phased vcf file as either ancestral or derived in a haplotype file and construct an associated genetic map

Using the phased data in the vcf file, Panderas converts ‘0’s and ‘1’s to nucleotides (e.g., A, C, G, or T) using the allele information in the ‘REF’ and ‘ALT’ columns, respectively, Then, these alleles are recoded as either ancestral or derived by comparing alleles at genomic coordinates in the vcf file to ancestral alleles at the same coordinates in the Homo_sapiens_hg38_reference file from Ensembl (https://ftp.ensembl.org/pub/release-112/fasta/ancestral_alleles/). This file contains high–confidence calls of ancestral state that have been confirmed by sequence data from different non-human primates (https://useast.ensembl.org/info/genome/compara/mlss.html?mlss=2006). When a given allele at a genomic coordinate in the vcf file matches the high-confidence ancestral allele at the same coordinate in the Homo_sapiens_hg38_reference file, this allele is coded as ‘0’ (for ancestral), and the non-ancestral allele at the same coordinate in the dataset is coded as ‘1’ (for derived). This process is applied to variation along a given chromosome, resulting in the recoding of ‘0’s and ‘1’s across individuals at each genomic coordinate. Alleles that are not coded as ‘0’ or ‘1’ are removed from the dataset.

To construct the associated genetic map, the Panderas software performs a linear interpolation to determine the genetic position (centimorgans, cM) of loci in a given dataset based on the physical position (i.e., genomic coordinate) and the corresponding genetic position in a generic PLINK map file (i.e., the reference). Specifically, Panderas searches for the two physical and genetic positions in the PLINK reference file that flank a query locus in a given dataset and calculates the genetic position for that query locus using information in the reference. Next, the Panderas software generates two formatted files, *.hap (containing the recoded phased alleles) and *.map (containing chromosome number, rs identifiers, genetic position and physical position), that serve as input files for selscan and HaploSweep (Szpiech and Hernandez 2014; Zhao *et al*. 2024)(**Fig. 1**). If users wish to include a population-specific genetic map instead, they need to replace the generic PLINK map with the population-specific genetic map and allow Panderas to create the associated map.

#### 2.1.3 Step 3: Run the selscan and/or HaploSweep programs using the input files generated by Panderas

The *.hap and *.map files, generated by Panderas, can be used directly as input for selscan and HaploSweep (Szpiech and Hernandez 2014; Zhao *et al*. 2024). More specific information on the usage of selscan and HaploSweep can be found at https://github.com/szpiech/selscan and https://github.com/ChenHuaLab/HaploSweep, respectively. The normalized output files from these programs will serve as input files for Panderas_plot in Step 3.

#### 2.1.4 Step 4: Plot the numerical output of selscan and/or HaploSweep with Panderas_Plots

For the *i*HS analysis, the normalized selscan and HaploSweep output files contain standardized *i*HS statistics that have either a negative or positive sign. Extreme negative standardized values indicate extensive haplotype structure around the ancestral allele whereas extreme positive standardized values denote unusually long haplotypes surrounding the derived allele. Users should be aware that selscan and HaploSweep by convention report positive statistics when the derived allele is under selection and negative statistics when the ancestral allele is favored (Szpiech and Hernandez 2014; Villegas-Mirón *et al*. 2021), which is opposite to the orientation described in Voight et al. (Voight *et al*. 2006).

The Panderas_Plots software also offers an array of options to visualize and annotate the tabular output of selscan and HaploSweep (**Fig. 1**). For example, Panderas_Plots can generate Manhattan plots of standardized statistics and insert a user-specified dashed horizontal line to indicate extreme values (specifically, standardized statistics above this threshold line will be considered outliers; **Fig. 1B**). Other noteworthy features include: i) annotating specific *i*HS values with their corresponding SNP identifiers using the --snps–to-highlight flag; ii) changing the size and/or color of each dot in Manhattan plots with the --size–dots flag; and iii) creating a list of genes harboring outlier *i*HS values in a *.txt file using --gene– annot and --gene-file. A more complete list of features can be viewed using the “--help” flag as described in the README file in our GitHub repository (https://github.com/alisi1989/Polaris). Users also have the option to save the Manhattan plots as *.pdf, *.eps, *.svg, and/or *.png files.

Furthermore, Panderas_Plots can create EHH line graphs showing the decay of haplotype homozygosity on chromosomes around ancestral and derived alleles at the same core site (**Fig. 1C**). In addition, Panderas_Plots automatically distinguishes these distinct chromosomes by color (specifically, red color for chromosomes with the derived allele and blue color for chromosomes carrying the ancestral allele) and provide a corresponding legend (**Fig. 1C**). Like the Manhattan plots, the EHH colored graphs can be saved as *.pdf, *.eps, *.svg, and/or *.png files.

## 3 Results and discussion

To our knowledge, there are currently no publicly available methods that i) polarize alleles in phased data; ii) create associated genetic maps for selscan and/or HaploSweep analysis; and iii) plot the quantitative output. To demonstrate the utility of our approach, we applied the Panderas software to allelic variation on Chromosome 2. More explicitly, alleles along the entire length of the chromosome were recoded as either ancestral or derived in the Finnish population from the 1000 Genomes Project (The 1000 Genomes Project Consortium 2012) for downstream analysis by selscan and HaploSweep. Next, we used the Panderas_Plots software to visualize the numerical output of these analyses. We first focused our attention on the well-studied regulatory region in intron 13 of the *MCM6* gene, which contains single nucleotide polymorphisms associated with lactase persistence (Enattah *et al*. 2002, Ingram *et al*. 2007, 2009, Tishkoff *et al*. 2007, Hassan *et al*. 2016, Anguita-Ruiz *et al*. 2020, Campbell and Ranciaro 2021) and then examined other signals of selection across Chromosome 2. Our selscan and HaploSweep analyses both uncovered long-range haplotype homozygosity, spanning up to 2 million base pairs (Mb), around the functional derived T_−13910_ allele (rs4988235; *i*HS > 3.5) in the Finnish (**Table S1**; **Fig. 1A** and **1B, Figs. S1** and **S2**), indicative of a classic selective sweep. Prior studies also have reported similar genetic patterns in this population (Enattah *et al*. 2002; Poulter *et al*. 2003; Mulcare *et al*. 2004; Cramp *et al*. 2014; Campbell and Ranciaro 2021). Notably, we were able to immediately identify the target allele under selection based on the sign of the *i*HS statistic at rs4988235.

Furthermore, we examined both negative and positive outlier *i*HS statistics (*i*HS< -2 and *i*HS >2, representing the most extreme 5% of empirical values) on Chromosome 2 in the Finnish population (**Tables S2** and **S3**). More specifically, we compared the number of extreme negative and positive *i*HS statistics in two separate datasets: i) one consisting of polarized alleles generated with Panderas and ii) one containing unpolarized alleles in the original phased vcf file on Chromosome 2. Interestingly, this analysis revealed a significant excess of extreme negative and positive *i*HS statistics (p < 0.00001) in the unpolarized dataset (**Tables S2** and **S3**). Because our software classifies alleles based on high-confidence ancestral call in the Homo_sapiens_hg38_reference file, our polarized dataset is curated to include carefully characterized alleles. However, the loss of alleles due to this curation process (resulting in the polarized dataset) was only ∼8 % of the total number of alleles on Chromosome 2 (**Table S4**) and thus cannot account for the sizeable excess of outlier statistics in the unpolarized dataset. Taken together, these findings arguably suggest the possibility that a subset of the ‘hits’ of selection on Chromosome 2 in this dataset may be false positives.

Moreover, of the negative outlier *i*HS statistics in the unpolarized dataset which indicate selection on standing variation, both ancestral and derived alleles were coded as ‘0’ in the unpolarized dataset; likewise, a mixture of ancestral and derived alleles were coded as ‘1’ in the same dataset (**Tables S5** and **S6**). We also observed a similar pattern for the positive outlier *i*HS statistics in the unpolarized dataset (**Tables S5** and **S6**). Therefore, the positive or negative sign associated with these *i*HS statistics is not informative for identifying the mode of selection (i.e., either a classic or soft selective sweep, respectively) after applying the haplotype-based methods to unpolarized alleles.

In summary, the Polaris package has the capacity to efficiently classify alleles as either ancestral or derived along chromosomes and generate associated genetic maps. Our software also provides a range of option for visualizing signals of selective sweeps. While these strengths make Polaris a powerful tool for investigating, interpreting, and summarizing patterns of selection, it is also important to note the limitations of our software. Because we polarize alleles in a given dataset against high–confidence calls of ancestral state, alleles that match to a low-confidence ancestral allele in the dataset are subsequently removed. Although the loss of alleles will be low for many chromosomes, there are some chromosomes (e.g., 19, 21, and 22) that have a higher proportion of low-confidence ancestral alleles (∼25% to 43%; **Table S4**), leading to the exclusion of the corresponding alleles in the *.hap file. While the proportions of low-confidence ancestral alleles on these chromosomes seem high, the successfully polarized alleles that remain in the *.hap file can be considered highly reliable. In addition, although strongly negative *i*HS statistics could be indicative of selection on ancestral variation in a polarized dataset, it is possible that ancestral alleles in close proximity to a derived allele under selection could exhibit large negative values due to genetic hitchhiking and may not be the direct target of selection. Therefore, like with any analysis, it is important to corroborate signals of selection from these haplotype-based methods with additional lines of evidence. Nonetheless, if prudently applied, the Polaris package is highly beneficial for investigating the nature of selection on chromosomes and its role in human adaptation, providing new insights into the recent evolutionary history of modern humans.

## Acknowledgements

We thank the Center for Advanced Research Computing (CARC) at the University of Southern California for providing the computational resources for this project. We also thank our novice and experienced users for testing the reproducibility of our computational pipelines.

## Supplementary data

Supplementary data are available at *Bioinformatics* online.

## Data availability

The data underlying this article are available in the 1000 Genomes Consortium Project at https://www.internationalgenome.org/data-portal/population/FIN.

