## Supplementary Data for "Polaris: Polarization of ancestral and derived polymorphic alleles for inferences of extended haplotype homozygosity in human populations"

### Supplementary Tables

**Table S1—Truncated output from selscan and HaploSweep analyses**. This Table shows the truncated normalized output of the selscan and HaploSweep, focusing on the rs4988235 (C>T) single nucleotide polymorphism in intron 13 of the *MCM6* gene. Here, we provide the locus ID, chromosome number, physical position (genomic coordinate), allele frequency of the derived T-allele at rs4988235, the *i*HS statistic for rs4988235, and the method used to calculate the statistic.

**Table S2— Distribution of extreme positive and negative *i*HS statistics (*i*HS < -2 and *i*HS > 2) on Chromosome 2 in the Finnish population from the 1000 Genomes Project calculated by selscan**. We compared the number of extreme negative and positive *i*HS statistics—calculated by selscan (Szpiech and Hernandez 2014) —in two distinct datasets: i) one containing polarized alleles generated with Panderas and ii) one containing unpolarized alleles in the original phased vcf file on Chromosome 2 in a 2 × 2 contingency table. The chi-square statistic ($\chi^2$) statistic was used to measure the divergence between observed and expected data. Statistical significance was assessed at $P < 0.05$, which represents the probability of the observed data occurring by chance alone.

**Table S3—Distribution of extreme positive and negative *i*HS statistics (*i*HS < -2 and *i*HS > 2) on Chromosome 2 in the Finnish population from the 1000 Genomes Project calculated by HaploSweep**. We compared the number of extreme negative and positive *i*HS statistics—calculated by HaploSweep (Zhao et al. 2024) —in two separate datasets: i) one containing polarized alleles (generated with Panderas) and ii) one containing unpolarized alleles in the original phased vcf file on Chromosome 2 in a 2 × 2 contingency table. The chi-square statistic ($\chi^2$) statistic was used to measure the divergence between observed and expected data. Like in Table S2, statistical significance was assessed at $P <$ 0.05.

**Table S4—Proportion of alleles that are not classified as either ancestral or derived across the autosomal genome.** This Table shows the total number of high-confidence and low-confidence ancestral alleles per autosomal chromosome (this count does not include gaps, which indicate the absence of a particular allele in humans) in the Homo_sapiens_hg38_reference file. Furthermore, we show the number of low-confidence ancestral alleles (not including gaps) and the proportion of low-confidence ancestral alleles per chromosome. Importantly, low-confidence alleles are not used to categorize alleles as ancestral or derived in a given dataset.

**Table S5—Summary statistics for unpolarized SNPs on Chromosome 2 in the Finnish population from the 1000 Genomes Project calculated by selscan**. Here, we examined both negative and positive outlier *i*HS statistics (*i*HS < -2 and *i*HS > 2) calculated by selscan (Szpiech and Hernandez 2014), representing the most extreme 5% of empirical values on Chromosome 2. Of these negative outlier statistics indicating selection on standing variation in the unpolarized dataset, the alleles coded as ‘0’ are a mix of ancestral and derived alleles (as opposed to one allele state only). Similarly, of the positive outlier *i*HS statistics indicating a classic selective sweep, the alleles coded as ‘1’ in the same dataset are a mixture of ancestral and derived alleles. The proportion of ancestral alleles coded as ‘1’ is ~10%, while the proportion of derived alleles coded as ‘0’ is ~70%.

**Table S6—Summary statistics for unpolarized SNPs on Chromosome 2 in the Finnish population from the 1000 Genomes Project calculated by HaploSweep.** Here, we examined both negative and positive outlier *i*HS statistics (*i*HS < -2 and *i*HS > 2) calculated by HaploSweep (Zhao et al. 2024), representing the most extreme 5% of empirical values on Chromosome 2. Of these negative outlier statistics indicative of selection on standing variation in the unpolarized dataset, the alleles coded as ‘0’ are a mix of ancestral and derived alleles (as opposed to one allele state only). Similarly, of the positive outlier *i*HS statistics indicative of a classic selective sweep, the alleles coded as ‘1’ in the same dataset are a mixture of ancestral and derived alleles. The proportion of ancestral alleles coded as ‘1’ is ~23%, while the proportion of derived alleles coded as ‘0’ is ~59%.

**Supplementary Figure**

**Figure S1: Plots of iHS and EHH statistics on Chromosome 2 in the Finnish population from the 1000 Genomes Project**. **Panel A** shows a Manhattan plot of *i*HS statistics calculated by HaploSweep (Zhao et al. 2024). The dashed horizontal line indicates the threshold for outlier *i*HS statistics as specified by the user (statistics above this line are considered outliers). Here, we highlighted the derived $T_{-13910}$ allele associated with lactase persistence in the Finnish population with a red dot and its corresponding rs identifier (rs4988235). The green and other red dots denote extreme *i*HS statistics. More explicitly, *i*HS > 4 (red dots) and *i*HS < -4 (green dots) indicate selection on derived alleles (suggestive of a classic selective sweep) and selection on standing variation (suggestive of a soft selective sweep). **Panel B** shows the decay of EHH (haplotype homozygosity) with increasing distance from the rs4988235 core site, which is associated with lactase persistence. In this EHH line graph, the x-axis shows physical position distance on Chromosome 2 in megabases (Mb), and the y-axis is the probability that two chromosomes are homozygous at all SNPs for the interval from the core site to distance x. Lastly, the blue line represents the decay of homozygosity of chromosomes carrying the ancestral allele at the core, while the red line signifies the decay of homozygosity on chromosomes carrying the derived allele at the core site.

**Table S1—Truncated output from selscan and HaploSweep analyses**

| locusID | chr | Genomic coordinate | Frequency of derived allele | iHS statistic | Method |
|---|---|---|---|---|---|
| rs4988235 | 2 | 135851076 | 0.590909 | 3.80086 | selscan |
| rs4988235 | 2 | 135851076 | 0.590909 | 3.68285 | HaploSweep |

**Table S2—Distribution of extreme positive and negative *i*HS statistics (iHS < -2 and iHS > 2) calculated by selscan on Chromosome 2 in the Finnish population from the 1000 Genomes Project.**

| | Positive *i*HS | Negative *i*HS | Total |
|---|---|---|---|
| **Polarized dataset** | 9597 | 10821 | 20418 |
| **Unpolarized dataset** | 10375 | 13220 | 23595 |
| **Total** | **19972** | **24041** | **44013** |

$X^2$ = 40.5831, df=1, $P < 0.00001$ (two-tailed *P*-value)

**Table S3—Distribution of extreme positive and negative *i*HS statistics (iHS < -2 and iHS > 2) calculated by HaploSweep on Chromosome 2 in the Finnish population from the 1000 Genomes Project**.

| | Positive *i*HS | Negative *i*HS | Total |
|---|---|---|---|
| **Polarized dataset** | 9390 | 10650 | 20040 |
| **Unpolarized dataset** | 11399 | 11050 | 22449 |
| **Total** | **20789** | **21700** | **42489** |

$X^2$ = 65.1446, df=1, $P < 0.00001$ (two-tailed *P*-value)

**Table S4—Proportion of alleles that are not classified as either ancestral or derived across the autosomal genome.**

| Chromosome number | Total number of alleles | Total number of low-confidence alleles | Proportion of unclassified alleles |
|---|---|---|---|
| 1 | 244,617,005 | 42,195,906 | 0.1724978 |
| 2 | 238,669,784 | 20,597,194 | 0.0850444 |
| 3 | 195,672,565 | 16,398,223 | 0.0838044 |
| 4 | 187,045,454 | 14,913,231 | 0.0797305 |
| 5 | 178,770,047 | 21,120,058 | 0.1181409 |
| 6 | 168,120,072 | 14,518,714 | 0.0863592 |
| 7 | 156,160,701 | 21,540,850 | 0.1379403 |
| 8 | 142,898,027 | 12,977,994 | 0.0908200 |
| 9 | 136,184,977 | 33,447,690 | 0.2456048 |

| | | | |
|---|---|---|---|
| 10 | 131,810,276 | 14,722,457 | 0.1116943 |
| 11 | 132,125,409 | 16,486,958 | 0.1247826 |
| 12 | 131,156,653 | 12,413,281 | 0.0946447 |
| 13 | 112,823,488 | 25,604,559 | 0.2269435 |
| 14 | 105,496,039 | 25,438,373 | 0.2411311 |
| 15 | 100,550,156 | 29,038,865 | 0.2887998 |
| 16 | 88,553,477 | 25,484,395 | 0.2877854 |
| 17 | 81,340,382 | 14,835,543 | 0.1823884 |
| 18 | 79,248.572 | 10,682,179 | 0.1347933 |
| 19 | 55,880,343 | 14,212,706 | 0.2543418 |
| 20 | 63,587,922 | 9,107,578 | 0.1432281 |
| 21 | 46,070,973 | 15,090,765 | 0.3275547 |
| 22 | 49,696,577 | 21,479,012 | 0.4322030 |

**Table S5—Summary statistics for unpolarized SNPs on Chromosome 2 in the Finnish population from the 1000 Genomes Project calculated by selscan**.

| Summary Statistic | Quantitative outcome |
|---|---|
| Total SNPs with \| iHS \| > 2 | 23,595 |
| Total SNPs with iHS > 2 | 10,375 |
| Total SNPs with iHS < -2 | 13,220 |
| Count of ancestral alleles (coded as ‘1’) in ALT column | 1,049 |
| Count of derived alleles (coded as ‘0’) in REF column | 9,276 |
| Proportion of ancestral alleles (coded as ‘1’) in ALT column | 0.1011 |
| Proportion of derived alleles (coded as ‘0’) in REF column | 0.7017 |

**Table S6—Summary statistics for unpolarized SNPs on Chromosome 2 in the Finnish population from the 1000 Genomes Project calculated by HaploSweep.**

| Summary Statistic | Quantitative outcome |
|---|---|
| Total SNPs with \| iHS \| > 2 | 22,449 |
| Total SNPs with iHS > 2 | 11,399 |
| Total SNPs with iHS < -2 | 11,050 |
| Count of ancestral alleles (coded as ‘1’) in ALT column | 2,677 |
| Count of derived alleles (coded as ‘0’) in REF column | 6,493 |

| | |
|---|---|
| Proportion of ancestral alleles (coded as ‘1’) in ALT column | 0.2348 |
| Proportion of derived alleles (coded as ‘0’) in REF column | 0.5876 |

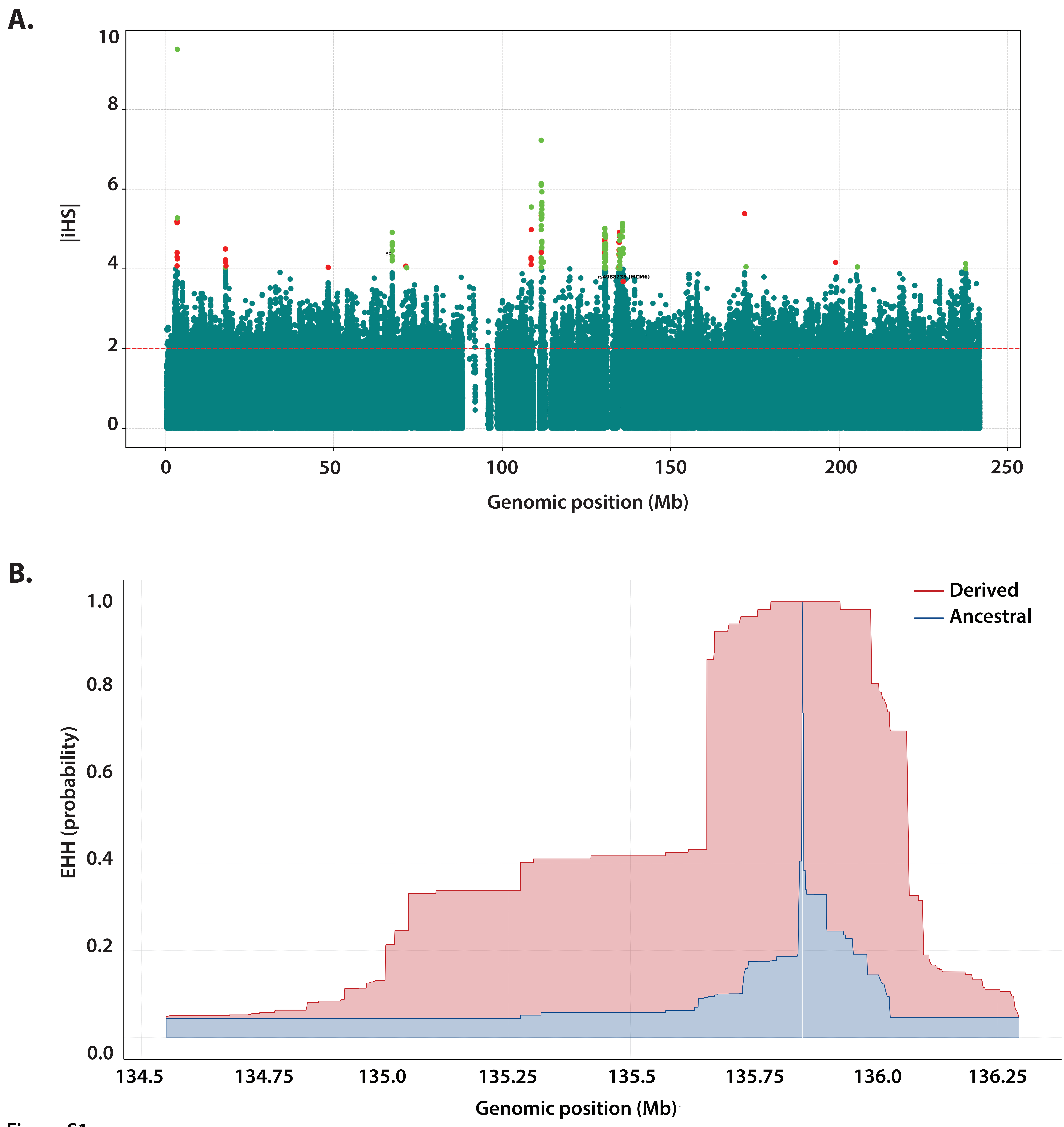
A.
10
8
6
4
2
0
|iHS|
rs4988235 (MCM6)
0
50
100
150
200
250
Genomic position (Mb)
B.
Derived
Ancestral
1.0
0.8
0.6
0.4
0.2
0.0
EHH (probability)
134.5
134.75
135.0
135.25
135.5
135.75
136.0
136.25
Genomic position (Mb)

Figure S1
